# Biofilm formation during pneumococcal carriage imprints naturally acquired humoral immunity

**DOI:** 10.64898/2025.12.17.694863

**Authors:** Jessica R. Lane, Henry Mauser, Silvia E. Santana-Krímskaya, Vahini S. Konda, Andrew DePass, Giuseppe Ercoli, Federico I. Prokopczuk, Mohammed Mohasin, Adonis D’Mello, Hervé Tettelin, Jeremy S. Brown, Luis F. Reyes, Carlos J. Orihuela

## Abstract

*Streptococcus pneumoniae* (Spn) colonization of the nasopharynx is a prerequisite for transmission and invasive disease. To investigate how repeated asymptomatic colonization shapes immunity and influences bacterial traits, we developed the Repeated Asymptomatic Murine Pneumococcal Colonization (RAMPC₃) model using strains belonging to serotypes: 2 (D39), 3 (WU2), and 4 (TIGR4). Sequential colonization revealed strain- and exposure-order–dependent effects on bacterial burden, with initial colonization yielding robust carriage and subsequent exposures resulting in diminished burden and rapid clearance. Humoral profiling demonstrated antigenic imprinting: the first colonizing strain largely determined IgG and IgA specificity, with minimal diversification after repeated exposures. Reactivity was strongest for biofilm-associated antigens correlating with each strain’s biofilm-forming capacity. Using TIGR4 mutants deficient in biofilm formation, we confirmed that in vivo aggregate formation drives humoral responses. Human sera from naturally colonized adults mirrored these findings, favoring biofilm antigens independent from capsule. Protection was demonstrated as triple-colonized mice exhibited reduced mortality and bacteremia following pneumococcal pneumonia challenge. Moreover, the initial colonizing strain influenced protection against heterologous infection, underscoring the lasting imprint of the biofilm phenotype on immunity. Finally, IgA responses in nasal-associated lymphoid tissue paralleled serum IgA patterns, validating systemic measurements as a proxy for mucosal immunity. Collectively, these results reveal that biofilm formation during colonization is a key determinant of humoral immunity and protection, providing insight into pneumococcal biology and informing strategies to design next-generation interventions.

**Author summary:** *Streptococcus pneumoniae* (Spn) is a leading cause of pneumonia, meningitis, and sepsis, yet its primary lifestyle is asymptomatic colonization of the nasopharynx. Understanding how colonization shapes immunity and bacterial physiology is critical for predicting disease risk and improving future vaccines. Using a novel murine model of repeated colonization and human sera from naturally colonized adults, we show that humoral immunity is strongly biased toward antigens expressed during biofilm growth, the predominant mode of *Spn* in the nasopharynx, rather than planktonic forms. This response is strain-dependent and imprinted by the first colonizing strain, influencing subsequent exposures and protection against pneumonia. Importantly, biofilm formation, not capsule, drives immune recognition revealing a key link between bacterial physiology and host immunity. These findings provide fundamental insight into pneumococcal biology and the host response, suggesting that targeting biofilm-associated antigens may improve vaccine design or strategies to prevent transmission and invasive disease.

## Introduction

*Streptococcus pneumoniae* (Spn), commonly known as the pneumococcus, is a Gram-positive bacterium and a major cause of otitis media, community-acquired pneumonia, bacteremia, and meningitis [1]. As a pathobiont that colonizes the nasopharynx, the risk of life-threatening infection is greatest among infants and the elderly, where host immunity is diminished [2–9]. Nasopharyngeal carriage is highly prevalent among young children, particularly in daycare settings, and serves as an important immunizing event [10–12]. Colonization induces both humoral and cellular adaptive immune responses against pneumococcal antigens including its capsule, pneumococcal surface protein A (PspA), its pore-forming toxin pneumolysin, and others. Repeated exposures are thought to strengthen these responses, reducing colonization burden and conferring protection against disease [13–16].

Naturally acquired immunity to *Spn* primarily arises from the host response to its protein antigens, whereas vaccine-induced immunity instead targets the bacterium’s exopolysaccharide capsule. *Spn* encompasses more than 100 biochemical and antigenically distinct serotypes [17–19], and therefore antibodies against one capsule type are unable to protect against infection caused by *Spn* carrying a different capsule type [20]. Instead, licensed vaccines achieve broad protection by being polyvalent, currently incorporating capsular polysaccharide (CPS) from up to 21 distinct serotypes. Importantly, the immunological steps by which *Spn* colonization primes protein-specific immunity remain poorly defined. This represents a critical gap in our understanding of pneumococcal host-pathogen interactions and an opportunity to improve the design of prophylactic measures.

*Spn* forms biofilms during nasopharyngeal colonization, which, when compared to the planktonic state, enhances adhesion to host surfaces, provides protection against host-derived antimicrobial molecules, and promotes survival on fomites [21–28]. In contrast, during pneumonia and invasive disease, pneumococci adopt a planktonic phenotype, i.e., growing as individual diplococci in suspension, a state that facilitates complement evasion [29–31]. These two growth modes are driven by environmental and physiological cues and are transcriptionally and phenotypically distinct [32–35]. This distinction is critical because most studies examining antigenic responses to *Spn* colonization have not taken into account the biofilm phenotype and instead relied primarily on planktonic cell lysates to identify immunogenic proteins [16, 36–38]. Consequently, gaps exist in our understanding of how biofilm-specific antigens contribute to naturally acquired immunity and protection against *Spn*.

In this study we utilize a novel murine model of repeated asymptomatic colonization to characterize the development of humoral immunity against *Spn* and assess the contribution of antigens produced during biofilm and planktonic growth. We evaluate how repeated colonization influences bacterial burden, the duration of carriage, and in turn, protection against lethal challenge by unrelated strains of *Spn*. Using human sera, we validate the immunogenicity of biofilm-associated antigens, confirming their potential as meaningful targets. Our findings provide insight into the dynamics of the immune response to *Spn* colonization, describe a striking outcome for early exposure, and provide important new considerations for the selection of protein antigens for a next-generation pneumococcal vaccine.

## Results

### *Spn* burden in a murine repeated asymptomatic colonization model depends on strain and exposure order

To characterize how the adaptive immune response develops following repeated exposures to *Spn*, we created the Repeated Asymptomatic Murine Pneumococcal Colonization (RAMPC₃) model using serotype 2 strain D39, serotype 3 strain WU2, and serotype 4 strain TIGR4. These strains represent distinct genetic lineages and are frequently used by *Spn* investigators. RAMPC₃ involves sequential, non-overlapping colonization of mice with these strains over a three-month period (Fig 1A). To dissect strain-level effects, two cohorts of mice were colonized with these strains but in opposite order: Cohort A (WU2 → D39 → TIGR4) and Cohort B (TIGR4 → D39 → WU2). Nasal wash (NW) bacterial burden was quantitated on post-inoculation days 1, 3, 7, and 14, and serum was collected on days −7, 21, 49, and 79. Colonization by each strain did not persist beyond 28 days.

**Fig 1.**
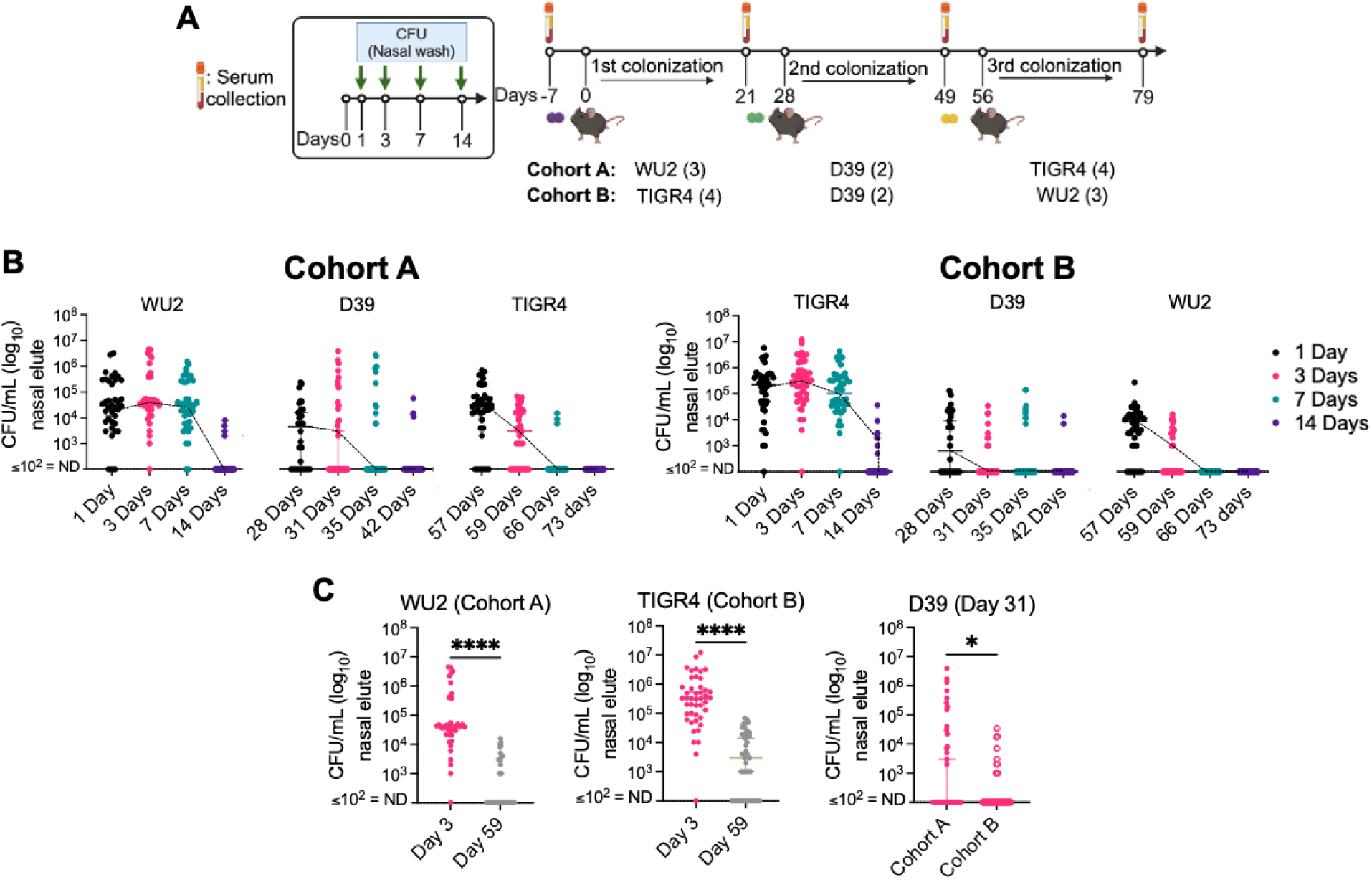
*Spn* burden in a murine repeated asymptomatic colonization model depends on strain and order exposure. **(A)** Schematic of the Repeated Asymptomatic Murine Pneumococcal Colonization (RAMPC_3_) model (see methods). 9-week-old C57BL/6J male and female mice were intranasally inoculated with 10^6^ CFU/mL of WU2 (Cohort A) or TIGR4 (Cohort B) for the first colonization event, followed by D39 for the second, and the final and third event with TIGR4 or WU2, respectively. **(B)** Bacterial burden was determined over a 2-week period post-inoculation by colony forming units (CFUs) obtained from nasal washes with saline. **(C)** Bacterial burden for each strain at Day 3 and Day 59 or Day 31. Each dot is one mouse sample. N=40-45 per group. Not detected (ND) ≤ 10^2^ CFU/mL. Mann-Whitney t-test and median with 95% confidence interval (CI). *=p ≤0.0332; ****=p≤ 0.0001.

Our first observation was that initial colonization, regardless of the strain used, resulted in robust carriage that remained stable for at least seven days. For Cohort A, median bacterial titers in NW of mice colonized with WU2 exceeded 10^4^ CFU/mL for the first 7 days whereas for Cohort B, median bacterial titers in NW for TIGR4 exceeded 10^5^ CFU/mL over the same time. Second exposures, both with D39, resulted in median initial burdens that were 10-fold and 100-fold lower than for WU2 or TIGR4, respectively, with most animals having cleared D39 within 7 days of challenge (corresponding to day 35; Fig 1B). For WU2 and TIGR4, median bacterial burden in NW differed by more than 100-fold depending on whether the strain was first or third in the corresponding cohort (Fig 1C). Initial D39 burden in RAMPC_3_ was consistently lower than in single-strain colonization (S1 Fig). Overall, our results with RAMPC_3_ align with human epidemiology [13–16], i.e., repeated colonization can occur but coincides with diminished burden and more rapid clearance. This decline likely reflects the development of adaptive immunity.

### The first colonization event imprints a humoral immune response to pneumococcal proteins consistent throughout repeated colonization

Using RAMPC_3_ serum samples, we examined how humoral immunity developed over time. Notably, and instead of meaningfully increasing the diversity of antigens recognized, immunoblots using pneumococcal whole cell lysates (WCL) and sequential serum samples from the same mice revealed minimal acquisition of new IgG-reactive protein bands after the second and third colonization events, regardless of cohort (Fig 2A and 2B). Though sera from both cohorts on average detected 6-7 proteins in our WCL panel, strain-specific effects were evident: Cohort A showed a broader range of protein recognition, up to 20 bands, compared to Cohort B, up to 9 bands. A similar pattern was observed for IgA, albeit less pronounced (S2A and S2B Fig). These findings suggest antigenic imprinting had occurred, whereby the first colonizing *Spn* strain encountered largely determines the humoral immune response to repeated colonization events.

**Fig 2.**
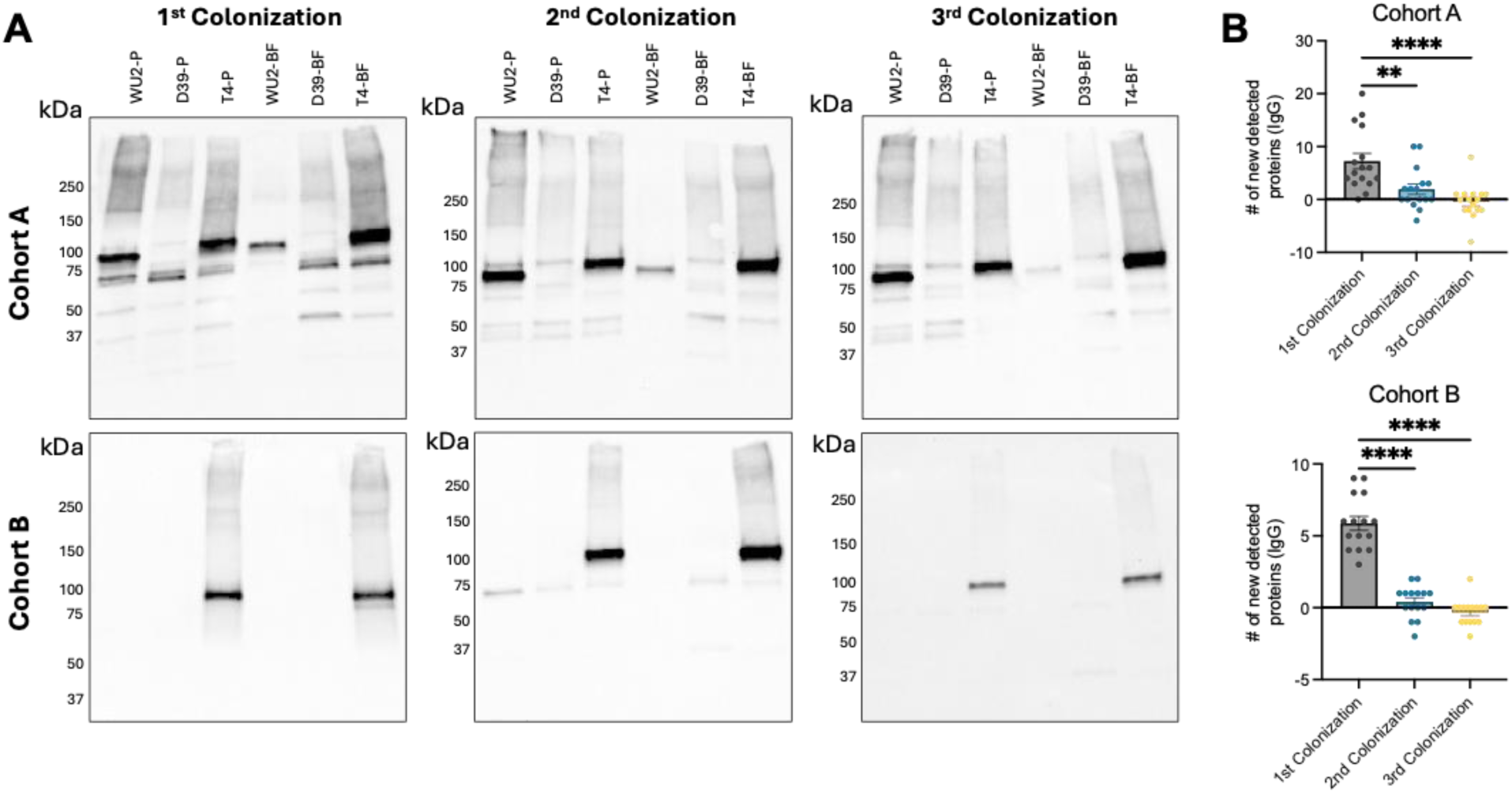
The first colonization event imprints a mucosal and systemic antibody response to the proteins that persist following repeated colonization. **(A)** Equal amounts of whole bacterial cell lysates grown planktonically (P) or in a biofilm (BF) from three *Spn* strains WU2 (serotype 3), D39 (serotype 2), and TIGR4 (serotype 4) were analyzed by immunoblot. Membranes were probed individually with mouse sera (1:1000) from Cohort A and Cohort B RAMPC_3_ mice after the first, second, and third colonization events and secondary α-mouse IgG (1:10000). Representative blots shown. **(B)** The number of new protein antigens detected by IgG on immunoblots from Cohort A and Cohort B RAMPC_3_ mice after the first, second, and third colonization events. N=15-16. One-way ANOVA and mean with standard deviation. **=p≤ 0.002; ****=p≤ 0.0001.

As the detected proteins are immunogenic and therefore viable targets for intervention, we probed a previously described *Spn* protein array using sera from both RAMPC_3_ cohorts after the first and third colonization events [39–41]. The six strongest signals included Pneumococcal histidine triad protein D (PhtD), Lysozyme M (LysM), Pneumococcal surface adhesin A (PsaA), Lytic transglycosylase G, a serine protease (Subtilase family-S8 subtilisin), and PspA (Fig 3A). These proteins play key roles in surface interactions of *Spn*, including adhesion and cell wall hydrolysis, and have been shown to be differentially expressed during colonization compared to bloodstream infection (Table 1) (S3 Fig). When stratified by cohort and colonization event, both cohorts of mouse sera consistently showed PhtD as the top hit followed by LysM, across all samples (Fig 3B). Notably, reactivity to individual proteins did not differ between the first and third colonization event (Fig 3B).

**Fig 3.**
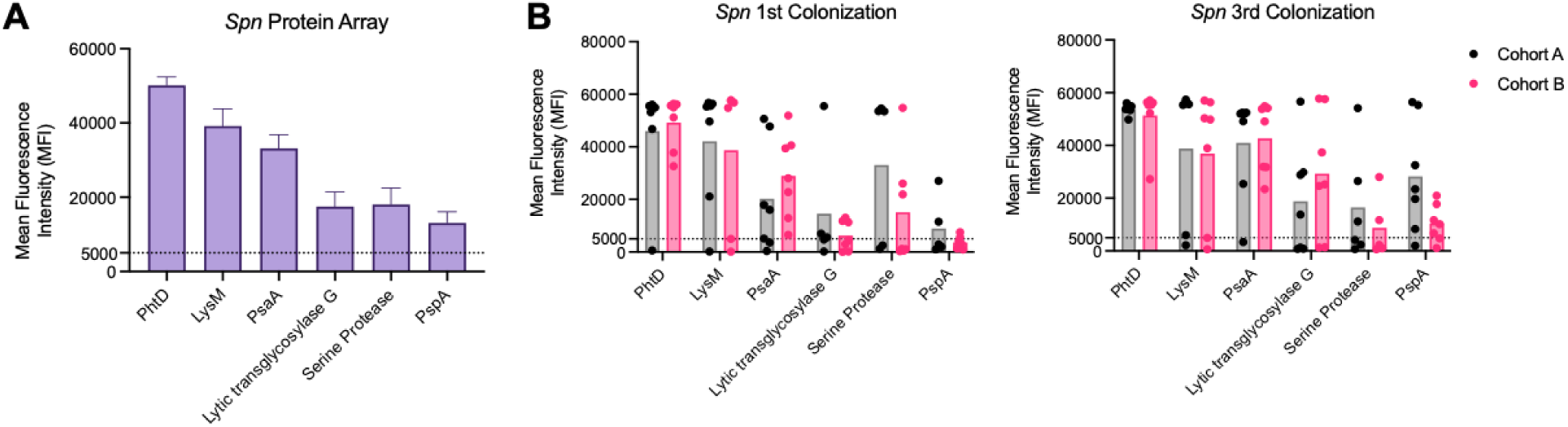
*Spn* protein array identifies specific antigens following repeated colonization. A pneumococcal protein array was constructed with 254 highly antigenic proteins. Proteins were selected from a panel of *Spn* strains and were conserved for recognition by IgG from healthy human adults (Croucher *et al.* 2017). Sera from RAMPC_3_ colonized mice after the first and third colonization events were used to probe the protein array (1:100) (see methods). **(A)** Both cohorts and timepoints combined. **(B)** Cohorts and timepoints expanded. N=14. Mean with standard deviation.

**Table 1.** Antigenic pneumococcal proteins from colonized mice.

| Antigen | Description | Protective (?) |
| --- | --- | --- |
| Pneumococcal histidine triad protein D (PhtD) (~93 kDa) | Surface adhesin | Sepsis model (subcutaneous) |
| Lysozyme M (LysM)-host (~17 kDa) | Hydrolysis of peptidoglycan cell wall and bacterial lysis at mucosal surfaces | Literature not found |
| Pneumococcal surface adhesin A (PsaA) (~35 kDa) | Surface adhesin, lipoprotein Mn<sup>2+</sup> transporter, protects against oxidative stress | Single carriage event (enhanced when combined with PspA), not protective in sepsis model (intraperitoneal) |
| Lytic transglycosylase G (~61 kDa) | Cell wall hydrolase | Target in combination with antibiotic treatment |
| Serine protease (Subtilase family-S8 subtilisin) (150-240 kDa) | Surface adhesion during colonization (unknown host target) | Literature not found |
| Pneumococcal surface protein A (PspA) (~67 kDa) | Choline-binding protein, surface adhesin, protects against C-reactive protein | Colonization, pneumonia, and sepsis models |

### Repeated murine *Spn* colonization elicits a strain-dependent humoral immune response associated with biofilm formation

Pneumococci in the nasopharynx exist in biofilms and this mode of growth has a distinct protein profile [42]. Notably, WU2, D39, and TIGR4 differ in their ability to form biofilms in vitro, with WU2 producing very little biomass and both D39 and TIGR4 capable of producing a robust biofilm after 24 hours of growth in a polystyrene well (Fig 4A). When we tested RAMPC₃ mouse sera collected after the third colonization event for IgG reactivity against planktonic WCL using ELISA we observed that reactivity was generally low across all strains (Fig 4B). In contrast, when tested against biofilm WCL, strain-specific IgG reactivity was observed with significantly greater responses for D39 and TIGR4 biofilm WCL compared to WU2 in both RAMPC_3_ cohorts (Fig 4C). Consistent with this, IgG reactivity of mouse sera to biofilm WCL positively correlated with the strain’s ability to form robust biofilms in vitro (Fig 4D). Similar patterns were observed for IgA (S5A-C Fig).

**Fig 4.**
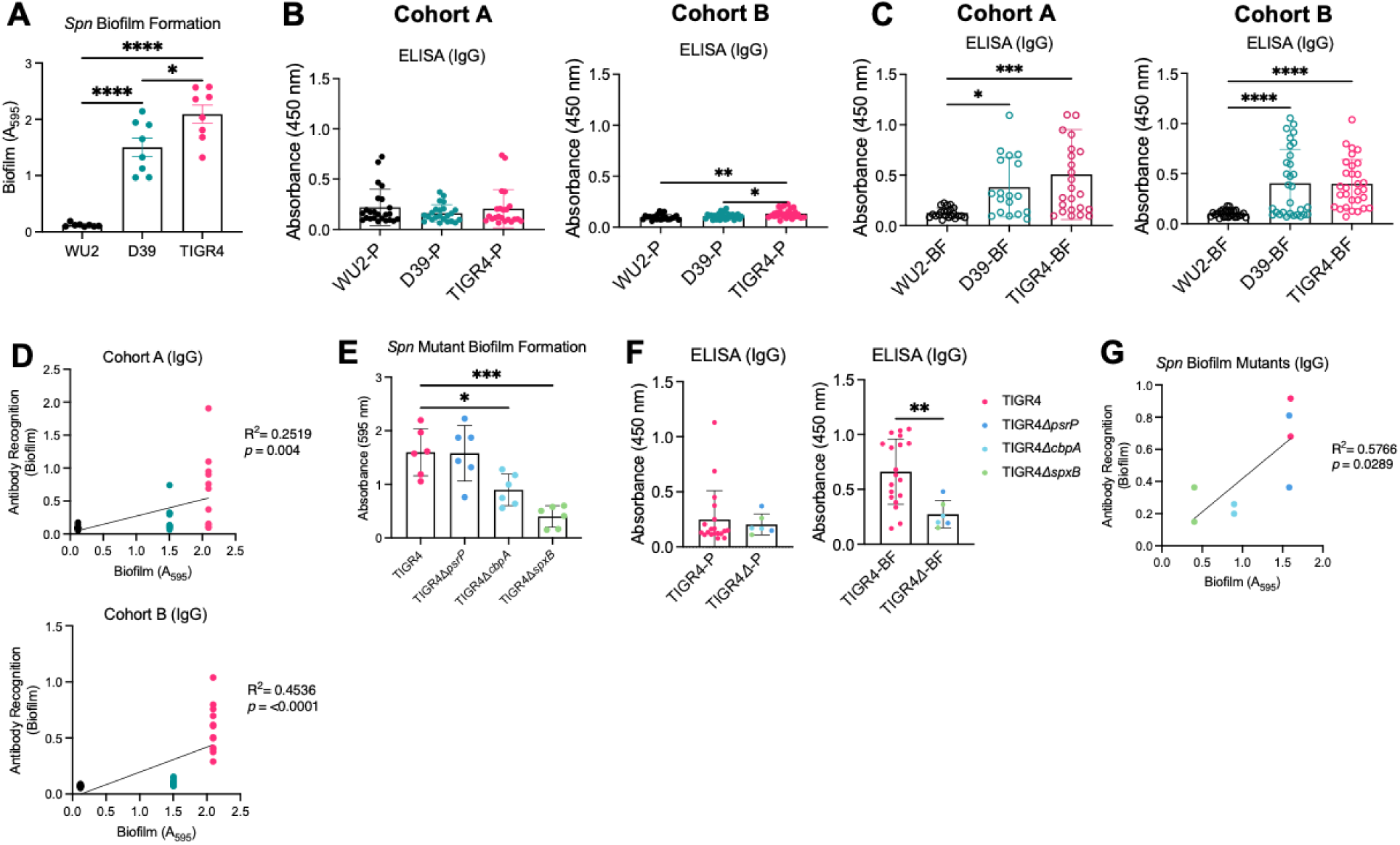
Repeated murine *Spn* colonization elicits a strain-dependent humoral response associated with biofilm formation. *Spn* lab strain’s (WU2, D39, TIGR4) ability to form biofilms as measured by crystal violet assay (see methods). N=8-10. Equal amounts of whole bacterial cell lysates grown **(B)** planktonically (P) or in a **(C)** biofilm (BF) from three *Spn* strains WU2 (serotype 3), D39 (serotype 2), and TIGR4 (serotype 4) were run on ELISAs and individually probed with serum (1:1000) from RAMPC_3_ mice in both Cohort A and Cohort B after the third colonization event. Secondary antibody α-mouse IgG (1:10000). Each dot is one mouse sample. N=24-30. **(D)** Linear regression correlation between ability of *Spn* strains to form biofilms and a ratio of antibody recognition to biofilm antigens. Each dot is one mouse sample. N=10-12. **(E)** Three isogenic TIGR4 mutant’s (Δ*psrP,* Δ*cbpA*, Δ*spxB*) ability to form biofilms as measured by crystal violet assay (see methods). N=6. **(F)** Equal amounts of whole bacterial cell lysates grown planktonically (P) or in a biofilm (BF) from TIGR4 and three isogenic TIGR4 mutants were run on ELISAs and individually probed with serum (1:1000) from mice colonized once with each strain for 21 days. TIGR4 mutants are pooled together. Secondary antibody α-mouse IgG (1:10000). Each dot is one mouse. N=2. **(G)** Linear regression correlation between ability of *Spn* TIGR4 mutant strains to form biofilms and a ratio of antibody recognition to biofilm antigens. One-way ANOVA or Mann-Whitney t-test and mean with standard deviation. *=p≤0.0332; **=p≤ 0.002; ***=p ≤0.0002; ****=p≤ 0.0001.

Given these results, we hypothesized that the ability of *Spn* to elicit a humoral response following asymptomatic colonization was dependent on its ability to form a biofilm in vivo. To test this, we colonized mice with a mixed panel of TIGR4 mutants previously shown to be attenuated in their ability to form in vivo aggregates on the surface of mucosal epithelial cells in the nasopharynx of mice during colonization (Fig. 4E and S5D Fig) [43, 44]. This panel included isogenic mutants deficient in *psrP* (encoding pneumococcal serine-rich repeat protein), *cbpA* (encoding choline-binding protein A), and *spxB* (encoding virulence factor pyruvate oxidase) [45, 46]. Notably, IgG reactivity of sera from these mice to planktonic WCL lysate was again low and equivalent to wildtype colonized mice whereas reactivity of mice colonized with the mutants to biofilm WCL was reduced (Fig 4F). We also saw a similar positive correlative effect with IgG recognition and biofilm formation ability as with the wildtype strains (4G) while IgA had a weaker signal (S5E Fig). Collectively, these results indicate that the strength of the humoral response to *Spn* varies in strain-dependent manner, is closely tied to the strain’s ability to form biofilms, and that biofilm formation *in vivo* is responsible for the strength of the humoral response that develops after initial colonization, with strain-specific consequence of this exposure influencing the response to and duration of repeated exposures.

### The human adult humoral immune response favors recognition of the biofilm phenotype and is not capsule-dependent

Given the importance of our findings, we sought to validate our results with human relevancy. We did so by evaluating IgA and IgG reactivity using sera from 17 human adults who were naturally and asymptomatically colonized with *Spn* at the time of collection. Similar to our observations in RAMPC_3_ mice, IgA and IgG reactivity to planktonic WCL was generally low; IgG responses to TIGR4 planktonic WCL were slightly stronger than those to WU2 or D39 (Fig 5A). Likewise, reactivity of IgA and IgG against biofilm WCL from D39 and TIGR4 was markedly higher than for WU2, with clear strain-specific differences. D39 elicited intermediate reactivity, whereas WU2 showed the weakest recognition (Fig 5B). This same pattern was observed when testing sera against WCL from *Spn* clinical isolates: IgA and IgG responses to planktonic WCL remained modest across all strains (Fig 5C), whereas IgG reactivity varied substantially for biofilm WCL, with distinct differences in recognition between strains (Fig 5D). Among these strains, 3-I (serotype 3, invasive) and 3-C (serotype 3, colonization) were the poorest biofilm producers, 4 was intermediate, and 2 produced the most biofilm biomass (Fig 5E). Accordingly, IgG reactivity also correlated positively with a strain’s ability to form biofilms in vitro (Fig 5F and 5G). For these experiments, sera were also probed against purified recombinant pneumolysin (rPly) and recombinant PspA (rPspA) as positive ELISA controls, both known to be strongly immunogenic [47–50]. Antibodies against these proteins detected an IgG response that favored rPspA over rPly (S6 Fig).

**Fig 5.**
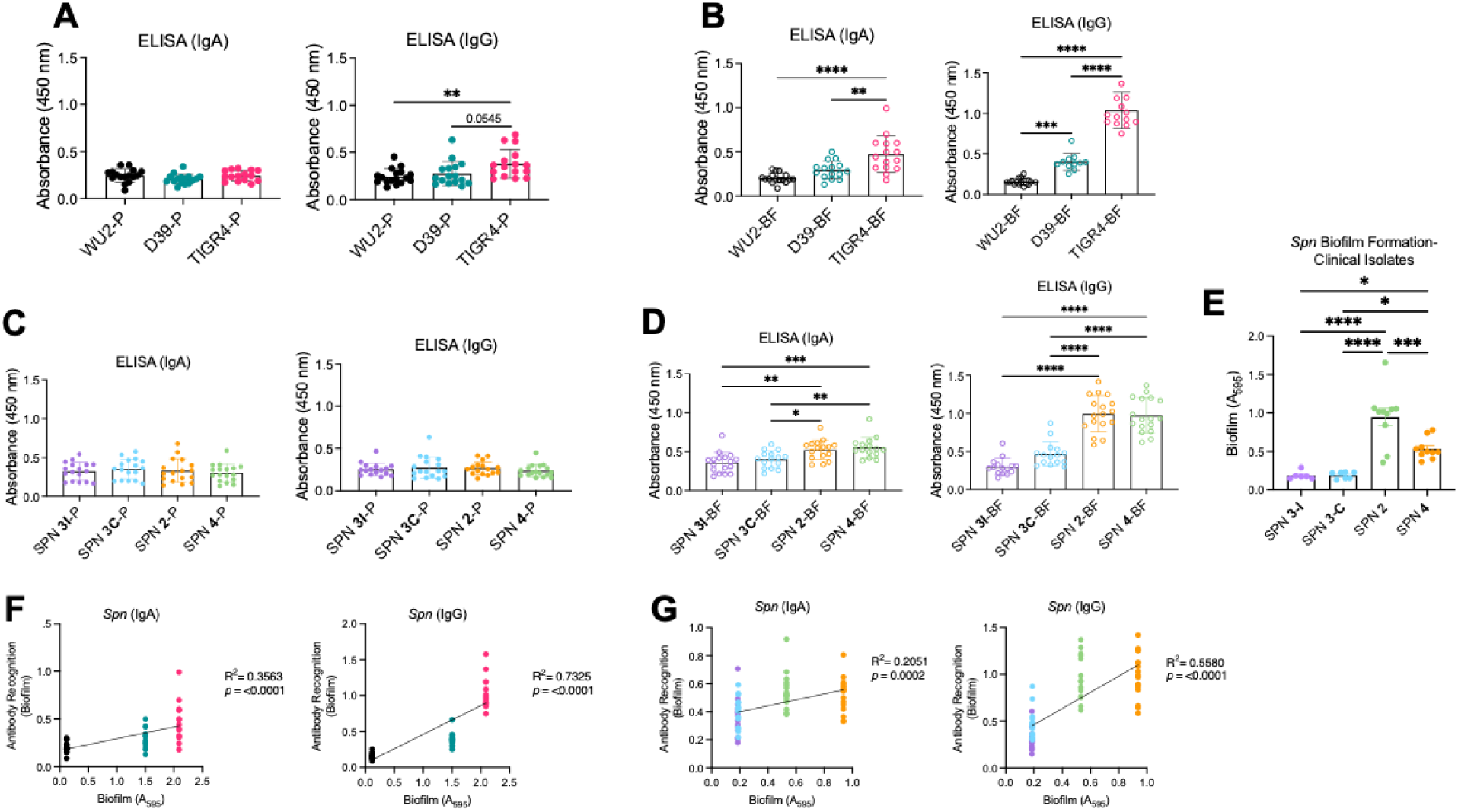
Serum antibodies from asymptomatic colonized human adults recognize *Spn* antigens depending on strain. Equal amounts of whole bacterial cell lysates (WCL) grown planktonically (P) or in a biofilm (BF) from **(A, B)** three *Spn* lab strains WU2 (serotype 3), D39 (serotype 2), and TIGR4 (serotype 4) and **(C, D)** corresponding serotype clinical isolates were run on ELISAs and individually probed with serum (1:1000) from asymptomatically colonized adults (aged 40-82) and secondary antibody α-human IgA and IgG (1:10000). Each dot is one human sample. N=17. **(E, F)** *Spn* lab strains (WU2, D39, TIGR4) and their corresponding clinical isolate’s ability to form biofilms as measured by crystal violet assay (see methods). N=8-10. **(G)** Linear regression correlation between ability of lab *Spn* strains and their **(H)** corresponding clinical isolates to form biofilms and antibody recognition to biofilm antigens. Mann-Whitney t-test, One-way ANOVA, and mean with standard deviation. *=p≤0.0332; **=p≤ 0.002; ***=p ≤0.0002; ****=p≤ 0.0001.

Two important considerations are that our ELISA-based results likely include reactivity to capsule present in WCL, and our human samples are confounded by vaccination and prior exposures to multiple and unknown-to-us *Spn* serotypes.

Therefore, it was important to determine the extent to which the observed responses were due to antibody against the capsular polysaccharide. To address this, we repeated the ELISA experiments using WCL from unencapsulated, i.e., rough (R), mutant derivatives of D39, WU2, and TIGR4. Importantly, IgA and IgG reactivity to planktonic WCL remained generally equivalent across strains, and we observed no difference in reactivity between encapsulated and unencapsulated WCL (S7A and S7B Fig).

Likewise, the response to biofilm WCL were again stronger overall and strain dependent. Notably, for WU2 and D39, the rough derivatives produced significantly more biofilm than their encapsulated counterparts (S7C Fig), but this effect only influenced IgA reactivity to D39. Overall, these results indicate that the preferential adult human humoral immune response to *Spn* biofilm WCL is independent of capsular polysaccharide.

### Repeated asymptomatic colonization with *Spn* protects against pneumococcal pneumonia

Finally, we tested whether triple-colonized mice were protected against pneumococcal pneumonia. To do this, mice were intratracheally infected with a fourth, unrelated *Spn* strain, 6A-10 (serotype 6A) [51], which is also forms a robust biofilm (Fig 6A). Mice in both Cohorts A and B exhibited reduced mortality compared to naïve controls, confirming that asymptomatic colonization can elicit protective immunity (Fig 6B). Consistent with this, both cohorts also showed a modest reduction in blood bacterial burden two days post-infection compared to controls (Fig 6C). No differences in IgA or IgG as measured by ELISA for 6A-10 WCL were observed between the cohorts (S8 Fig). We also sought to determine the extent to which the first colonizing strain influenced infection outcomes by colonizing mice with either WU2 or TIGR4, followed by intratracheal infection with the opposing strain or with D39. In this experiment, WU2 asymptomatic colonization protected mice against TIGR4 and D39 disease compared to naïve controls. In contrast, TIGR4 asymptomatic colonization provided protection only against WU2, with no significant differences in blood bacterial burden (Fig 7A-B and S9A and S9B Fig). Importantly, robust IgG responses and weaker IgA responses were again detected for biofilm antigens of the corresponding colonizing and infecting strains (Fig 7C-E and S9C-E Fig), providing further evidence of the critical impact of the initial colonizing strain and the lasting imprint of its biofilm phenotype on humoral immunity.

**Fig 6.**
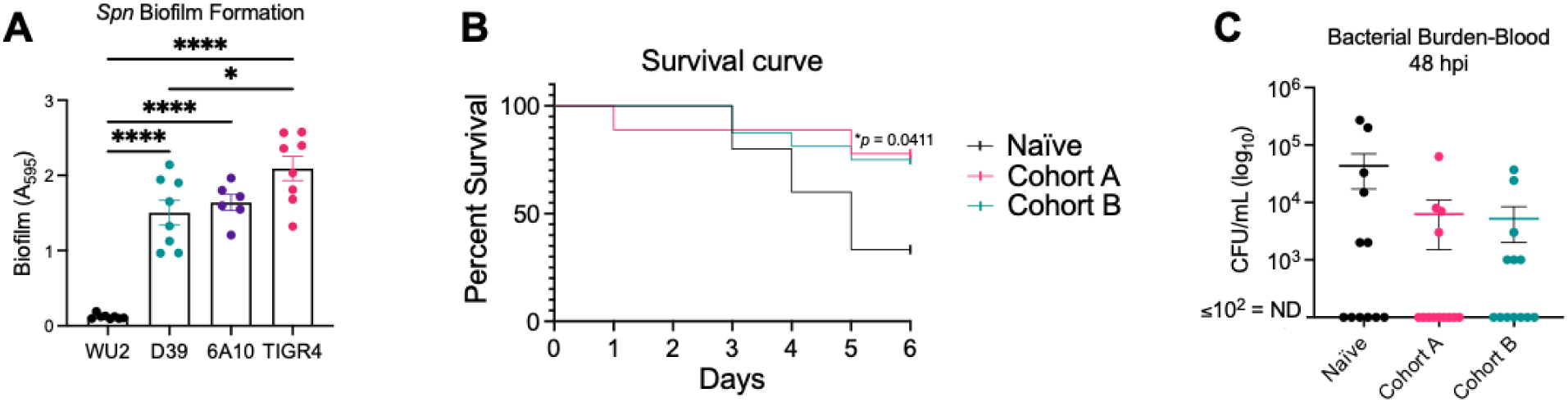
Repeated asymptomatic colonization with *Spn* protects against pneumococcal pneumonia. A cohort of age-matched female naïve mice and the RAMPC_3_ mice from both Cohorts A and B were intratracheally challenged with 10^5^ CFU/mL of *Spn* strain 6A-10 (serotype 6A) (see methods). **(A)** Biofilm formation for D39, WU2, TIGR4, and 6A-10 strains as determined by crystal violet assay. **(B)** Survival over time and **(C)** bacterial burden in the blood 48 hours post-infection (hpi) was recorded. Not detected (ND) ≤ 10^2^ CFU/mL. Kaplan-Meier survival curve with log-rank test. One-way ANOVA and mean with standard deviation or standard error of the mean (SEM). *=p≤0.0332; ****=p≤ 0.0001.

**Fig 7.**
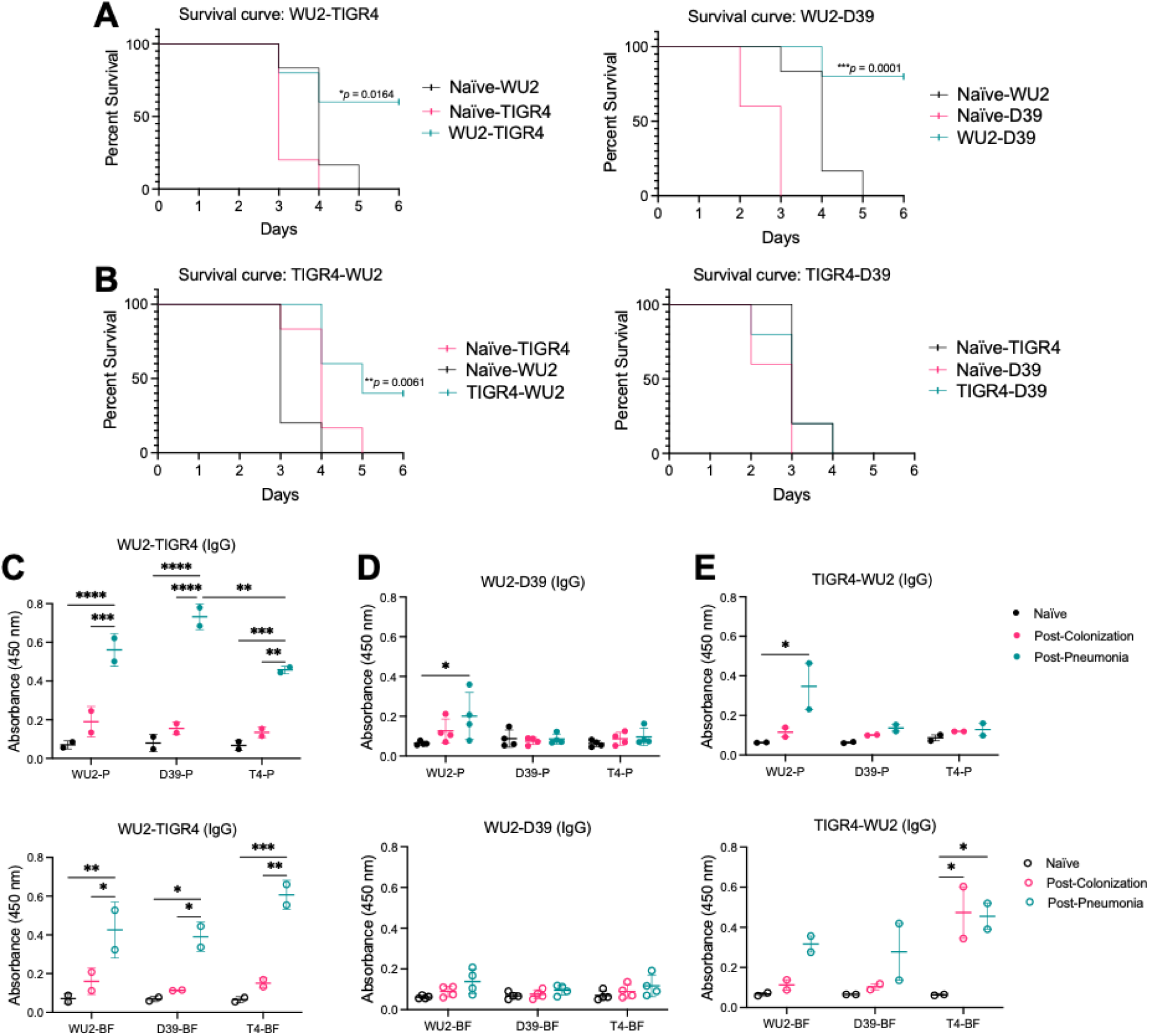
Protection against pneumococcal pneumonia is dependent on the colonizing *Spn* strain. 9-week-old C57BL/6J female mice were intranasally inoculated with 10^6^ CFU/mL of (A) WU2 or (B) TIGR4. After one month, mice were intratracheally challenged with 10^5^ CFU/mL of a different *Spn* strain: WU2 (serotype 3), D39 (serotype 2), and TIGR4 (serotype 4). Survival was recorded. N=5-6 per group. Kaplan-Meier survival curve with log-rank test. (C, D, and E) Equal amounts of whole bacterial cell lysates grown planktonically (P) or in a biofilm (BF) from three *Spn* strains (WU2, TIGR4, or D39) were run on ELISAs and were individually probed with mouse sera (1:1000) from surviving mice before and after colonization and infection and secondary α-mouse IgG (1:10000). Each dot is one mouse sample. N=2-4. Two-way ANOVA and mean with standard deviation. *=p≤0.0332; **=p≤ 0.002; ***=p ≤0.0002; ****=p≤ 0.0001.

### Serum IgA reactivity is comparable to mucosal tissue-derived antibody responses

Throughout our studies we had assessed IgA and IgG reactivity to pneumococcal antigens using serum. IgG is consistently and accurately measured using this method, providing a comprehensive view of the systemic humoral immune response during sequential colonization. This contrasts with IgA, the predominant mucosal-associated immunoglobulin found in airway secretions and tissue-specific sites such as nasal-associated lymphoid tissue (NALT) in mice or Waldeyer’s ring in humans [52]. To determine whether IgA derived from mucosal tissue mirrored our serum findings, we measured NALT IgA reactivity in three murine cohorts. Each cohort was colonized once with either WU2, D39, or TIGR4 and tested for reactivity against planktonic and biofilm cell lysates using ELISA. We observed similar patterns of IgA reactivity to biofilm bacterial lysates from NALT as those seen in serum (S10A-C Fig).

## Discussion

Recent conjugate vaccines targeting the *Spn* polysaccharide capsule have greatly reduced childhood incidence and mortality from pneumococcal infection. However, *Spn* remains a leading cause of community-acquired pneumonia, underscoring the need for improved protective strategies. Because colonization is a prerequisite for disease development, understanding the interactions *Spn* has with the host during colonization is critical for identifying or prioritizing antigens for next-generation protein-based vaccines. Our study demonstrates, using a novel repeated-colonization mouse model that was validated using human sera, that systemic humoral immunity strongly recognizes proteins produced preferentially during biofilm growth. Moreover, the first colonizing *Spn* strain imprints a stable pattern of antibody recognition that persists through subsequent colonization.

Over the past 25 years, *Spn* biofilm formation during colonization has gained recognition for its role in enhancing adherence to mucosal epithelial cells, protecting against desiccation, promoting survival on fomites, and facilitating bacterial interactions that increase competence and genetic exchange [23, 25, 27, 28, 53]. Although *Spn* in biofilms is less virulent than planktonic forms and transcriptomic studies reveal marked differences in gene expression between planktonic and biofilm states [35], biofilm bacteria can disperse under inflammatory conditions to cause disease [30, 43, 54].

However, most immunological studies have focused on planktonic antigens with their more commonly known impact on disease, overlooking that *Spn* typically exists as a biofilm during carriage. Our findings highlight that antibodies from experimentally colonized mice and asymptomatically colonized humans reacted strongly to biofilm-associated proteins, suggesting that this aspect of pneumococcal biology may provide a breakthrough in vaccinology.

Our findings with sera from colonized mice confirm that asymptomatic pneumococcal colonization induces a systemic humoral response to protein antigens. This was consistent with previous reports [41, 55] and aligns with controlled human infection studies in which subjects with higher antibody levels against pneumococcal proteins were better protected against rechallenge [56]. We did not identify specific proteins recognized by human sera due to unknown prior exposure history. However, antibodies in sera obtained from repeatedly colonized mice targeted surface adhesins (PhtD, PsaA, PspA, and a subtilisin-family serine protease) and cell wall hydrolases (LysM and lytic transglycosylase G). Cross-referencing these genes with in vivo RNA-seq data confirmed their expression was enhanced by *Spn* during nasopharyngeal colonization [35]. Many of these protein classes overlap with antigens recognized by IgG in other human serological studies of children and adults [57–59]. While proteins such as PspA and PsaA have been extensively characterized as protective antigens [60–66], to the best of our knowledge they have not been specifically evaluated for their ability to reduce colonization duration or bacterial burden during repeated colonization. Our findings suggest that such efforts are warranted and that the RAMPC_3_ model is well-suited to them.

Along such lines, we developed the RAMPC₃ model to create a multi-strain, multi-colonization system that better reflects human exposure while allowing control over strain order. RAMPC₃ findings showed that nasopharyngeal bacterial burden depended on both the number of colonization events and the strain order; these factors are critical for understanding heterologous cross-protection and disease prevention [67]. Our results with RAMPC_3_-generated sera mirrored that obtained with human sera, showing strong recognition of biofilm antigens, and further validating the model.

Because our laboratory strains were well characterized, we were able to link host response strength to each strain’s ability to form robust biofilms in vitro. We also had the ability to challenged mice with a fourth strain to assess the impact of colonization on infection and confirmed that IgA responses in NALT reflected those observed in serum. Herein, we used RAMPC_3_ to study the impact of repeated pneumococcal colonization on humoral immunity, however, the model is suitable for studying other aspects of colonization and disease progression.

We observed that robust biofilm-forming strains elicited stronger immune responses which correlated with reduced bacterial burden during subsequent colonization. However, protection against invasive disease after a single colonization event was not absolute and depending on the colonizing and infecting strain had varied outcomes following intratracheal challenge of mice. Given that we observed imprinting, which would limit the extent of humoral immunity which develops thereafter during repeated protection, we postulate that the observed gaps in protection are subsequently filled by the limited additional proteins whose recognition is added as well as contributions from other arms of the adaptative immune system, particularly IL-17 producing T-cells [15, 68]. Notably, the notion that biofilm antigens could be protective aligns with recent work by Jensen *et al.* who introduced the concept of biofilm-associated molecular patterns (BAMPs) as unique immunostimulatory molecules highly expressed in biofilms [69]. It remains important to consider that disease-causing pneumococci are likely planktonic and therefore not opsonized to the same extent as their biofilm counterparts by the colonization-induced humoral response, though the latter could still lower disease risk.

In conclusion, our study provides new insights into pneumococcal carriage and disease highlighting how biofilms influence humoral immunity and potentially other aspects of protection by using the novel triple colonization RAMPC_3_ model. Although the concept of “original antigenic sin” is well established in viral infection, its role in bacterial pathogenesis is not well appreciated with limited evidence of its occurrence or impact on outcomes regarding repeated infections. Our findings underscore that the biofilm phenotype plays a dominant role during colonization and is the primary version of the bacteria recognized by the humoral response against *Spn*. Collectively, this information has the potential to be leveraged regarding antigen selection for future vaccine formulations.

## Materials & methods

### Ethics statement

Human serum samples from adult volunteers aged 40-82 years were collected under Institutional Review Board (IRB) approved protocol at the Centro de Investigación Unisabana Center for Translational Science at the Universidad de La Sabana in Chia, Colombia (see Table S2). Serum samples were de-identified and did not meet the criteria for human subject research. Animal experiments were performed under the Institutional Animal Use and Care Committee (IACUC) approved protocol #22157 at The University of Alabama at Birmingham. Animal care and experimental protocols adhered to Public Law 89-544 (Animal Welfare Act) and its amendments, Public Health Services guidelines, and the Guide for the Care and Use of Laboratory Animals (U.S. Department of Health & Human Services).

### Bacterial strains

Four strains of *Spn* were used in this study (see Table S1). All bacteria were grown from frozen stock on tryptic soy agar plates with 5% sheep blood at 37°C and 5% CO_2_ overnight. Working broth cultures were grown in Todd-Hewitt broth with 0.5% yeast extract (THY) to exponential phase (OD_621_: ∼0.3-0.5) before being serially diluted in saline for experiments or harvested for protein. For 24-hour biofilm growth, a 1:100 culture of *Spn* was grown in THY in a sterile 100mm polystyrene plate at 37°C and 5% CO_2_ overnight.

### Bacterial lysis and protein quantification

For planktonic *Spn* lysates, 1 mL aliquots of bacterial cultures were spun down at ∼2,500xG for 3 minutes, then resuspended in protein lysis buffer (50mM Tris, 150mM NaCl, 1% TritonX-100) with 10% sodium dodecyl sulfate (SDS) and deoxycholate added in phosphate buffer saline (PBS). Lysates were incubated at 37°C and 5% CO_2_ until the liquid was clear. Biofilm *Spn* were washed twice with PBS after THY was removed, collected by washing with 1 mL of PBS, spun down and resuspended in protein lysis buffer, and then lysed as stated above. Protein concentration was quantified using the Pierce™ BCA Protein Assay Kit per the manufacturer’s recommendation (Thermo Scientific, #23225).

### Recombinant protein purification

Purification of recombinant proteins pneumolysin (rPly) and PspA (rPspA) was performed by using cobalt-affinity resin and the *E. coli* strains BL21(DE3) and NEBExpress® Iq carrying the respective plasmids [70, 71]. The strains were grown at 37°C in Luria-Bertani (LB) broth with required antibiotics, and expression was induced at OD_621_=0.4 by the addition of 1 mM isopropyl β-D-1-thiogalactopyranoside (IPTG).

After 4 hours of induction, cells were harvested and lysed in BugBuster Master Mix (MilliporeSigma, #71456-4) with 1 mM phenylmethylsulfonyl fluoride (PMSF) for 30 minutes at room temperature with gentle shaking. The cell lysates were spun at ∼3,000xG for 20 minutes, and the supernatant was loaded on beads pre-equilibrated with default buffer (5 mM Imidazole, 50 mM Tris-HCl pH 7.5, 150 mM NaCl). Beads were washed 10 times with the default buffer, and protein was eluted in the default buffer containing 100 mM imidazole. Collected protein fractions were confirmed on an SDS-PAGE gel and concentrated/buffer exchanged to PBS using a centrifugal concentrator tube. Protein concentration was measured by using the Pierce™ BCA Protein Assay Kit per the manufacturer’s recommendation. Equal loading of lysates was confirmed by Coomassie staining of SDS-PAGE gel and Ponceau staining of transferred proteins on nitrocellulose membrane.

### Immunoblots

Whole-cell bacterial lysates (WCL) (10 μg/mL) and recombinant protein (rPly: 0.1 μg/mL, rPspA: 1 μg/mL) were mixed with NuPAGE™ LDS Sample Buffer (4X) (ThermoFisher, NP0007) and β-mercaptoethanol before denaturing for 10 minutes at 95°C for 10 minutes. Samples were loaded onto an SDS-PAGE gel (4-15%; BioRad) before transferring onto a nitrocellulose membrane. After blocking with 5% bovine albumin serum (BSA), the membrane was incubated overnight at room temperature with the primary antibody, followed by washing with 1% tris-buffered saline-tween (TBS-Tween20) and incubation with the secondary antibody. Visualization of the blots was done using the Pierce™ ECL Western Blotting Substrate (ThermoFisher, PI32209) and ChemiDoc XRS+ System (BioRad). Primary antibody: human Colombia patient cohort (1:1000) or mouse serum (1:1000). Mouse NALT analysis (1:100). Secondary antibodies: HRP-conjugated goat α-human IgA (Invitrogen, #PA1-74395), HRP-conjugated goat α-human IgG (Jackson Immuno Research, #109-035-088), HRP-conjugated goat α-mouse IgA (Abcam, #ab97235), HRP-conjugate goat α-mouse IgG (Invitrogen, #31430) (all at 1:10,000).

### ELISA (enzyme-linked immunosorbent assay)

For detection of human and mouse IgA and IgG responses to various *Spn* lysates, a 96 well plate (ThermoFisher Immulon 4HBX, #3855) was coated with 2 μg/mL of rPly, rPspA, and the planktonic and biofilm-grown lysates of WU2, D39, TIGR4, and 6A-10 in PBS, covered, and placed at 4°C overnight. The next day, the plate was washed three times with PBS-Tween before blocking with 5% bovine albumin serum (BSA) in PBS for 1 hour at room temperature. The plate was washed twice more as above before the addition of desired primary serum diluted in PBS at (1:1000) and incubated for 3 hours at room temperature. After incubation with the primary serum, the plate was washed three times before incubation with the desired secondary antibody diluted in PBS at (1:10,000) and incubated for 1 hour at room temperature. The plate was washed five times before the addition of tetramethylbenzidine (TMB) substrate reagent (BD OptEIA TMB Substrate Reagent Kit, #555214) to each well-prepared according to the manufacturer’s recommendation and incubated in the dark for 10 minutes at room temperature. 2N sulfuric acid was added to each well to stop the reaction, and the absorbance was read on a BioTek Cytation 5 Cell plate reader at 450 nm. Primary antibody: human (Colombia patient cohort) or mouse sera (1:1000). Mouse NALT analysis (1:10). Secondary antibodies: HRP-conjugated goat α-human IgA (Invitrogen, #PA1-74395), HRP-conjugated goat α-human IgG (Jackson Immuno Research, #109-035-088), HRP-conjugated goat α-mouse IgA (Abcam, #ab97235), HRP-conjugated goat α-mouse IgG (Invitrogen, #31430) (1:10,000).

### Crystal violet assay for biofilm quantification

*Spn* lab and clinical strains of interest were grown as biofilms, as described above. After 24 hours, the media and planktonic bacteria were carefully aspirated, and the remaining biofilm layer was washed three times with PBS. The washed biofilms were allowed to dry completely before incubation with 0.1% crystal violet (Sigma-Aldrich, #C3886) stain in distilled water for 20 minutes at room temperature. The stain was removed, and the biofilms were washed as described. After completely drying again, the stained biofilms were resolubilized (95% ethanol and 5% acetic acid), and dilutions were added to a 96-well plate. The absorbance was read on a BioTek Cytation 5 Cell plate reader at 595 nm.

### Repeated Asymptomatic Murine Pneumococcal Colonization (RAMPC_3_) Model

C57BL/6J female and male mice aged ∼9 weeks in Cohort A were inoculated intranasally with *Spn* strain WU2 (∼10^6^ CFU/mL). At various timepoints (1, 3, 7, and 14 days), 10 μL of PBS was used to wash the nasal cavity of each mouse while under isoflurane sedation. ∼2 μL of recovered liquid was then diluted ten-fold, mixed thoroughly with a pipette, and plated onto blood agar plates for colony forming unit (CFU) enumeration. After ∼1 month, mice were inoculated with D39, and nasal washes were repeated as described. Another month later, mice were inoculated with TIGR4.

Serum from mice was collected via retro-orbital eye bleeds on days -7, 21, 49, and 79. Mice in Cohort B were colonized in the same manner, except the order of strains was TIGR4, D39, and WU2. Mice for NALT analysis were colonized once using the same method with D39, WU2, and TIGR4. NALT was harvested after ∼2 weeks, as described previously [72] and homogenized in 1 mL of PBS.

### Biofilm-deficient mutant colonization and ex vivo biofilm quantification

C57BL/6J female and male mice aged 8-10 weeks were inoculated intranasally as above with *Spn* strains TIGR4, TIGR4Δ*psrP*, TIGR4Δ*cbpA*, or TIGR4Δ*spxB* (10^6^ CFU/mL), with two mice per group. At 7 and 10 days, 10 µL of PBS was used to wash the nasal cavity of each mouse while under isoflurane sedation. ∼2 µL of recovered liquid was diluted ten-fold in PBS, and then 10 µL of the suspension was combined 1:1 with 1% crystal violet in H_2_O (Sigma-Aldrich, #C3886). 10 µL of this mixture was transferred to a glass slide, a coverslip was applied, and bacterial aggregates were counted at 1000x magnification using a Leica CME microscope. 50-100 aggregates were counted per sample and classified as 1, 2-9, or ≥10 diplococci. Number of aggregates for each of these three categories were expressed as a percentage of the total number of aggregates. On day 21 post-colonization, serum from mice was collected via retro-orbital eye bleeds.

### In vivo infections

C57BL/6J female and male mice aged ∼9 weeks or age-matched mice for the repeated colonization model were sedated using 2.5% vaporized isoflurane in oxygen. For intratracheal challenge (pneumonia model), the inoculum was prepared in PBS, and 100 μL containing ∼10^5^ CFU/mL was instilled into the lower airways by forced aspiration [73]. All animals were observed for recovery following infection and monitored daily afterward. Bacteremia was determined using blood obtained by tail bleed (∼2 μL of blood per mouse) diluted in PBS and enumerated via CFUs on blood agar plates. Serum was collected from surviving mice by retro-orbital eye bleeds.

### Pneumococcal protein array

A *Spn* protein array containing 254 proteins was constructed. Proteins were selected based on having a high level of conservation in a panel of >600 *Spn* strains and included the majority of the conserved proteins that were significantly recognized by IgG present in human sera obtained from healthy adults [58]. The array was constructed using genes amplified from bacterial genomic DNA (*Spn* strain TIGR4) and cloned into a T7 expression vector. Proteins were expressed by incubating the plasmids for 16 hours in *E. coli*-based *in vitro* transcription/translation (IVTT) reactions (RTS E. coli HY 100 kit, biotechrabbit, #BR1400106). Proteins were tested for expression by immunoblot using antibodies against N-terminal poly-histidine (His) after transfer onto nitrocellulose-coated glass AVID slides (Grace Bio-Labs, #305384), using an Omni Grid 100 microarray printer (Genomic Solutions). Arrays were probed with mouse serum samples diluted (1:100) in protein array blocking buffer (GVS, #10485356) and supplemented with *E. coli* lysate. Images were acquired using a Innoscan 710G scanner (Innopsys) and analyzed using Mapix 8.5 software. ‘‘No DNA’’ controls consisting of *E. coli* IVTT reactions without the addition of DNA were averaged and used to subtract background *E. coli* reactivity. All results presented are expressed as Mean Fluorescence Intensity (MFI).

### Statistical analysis

All statistical analyses were done using GraphPad Prism (version 10). For comparisons of two groups, the nonparametric two-tailed Mann-Whitney log-rank *t*-test was used, and the median with a 95% confidence interval or the standard deviation (SD) is shown. For all multiple comparisons, a nonparametric one-way or two-way analysis of variance (ANOVA) with Tukey’s pairwise test was used, and the SD is shown.

## Acknowledgements

This work was supported by National Institutes of Health (NIH) T32 grant 2T32AI007051-46 (JRL), NIH F31 grant 1F31AI186511-01 (JRL), NIH R01 grant 5R01AI156898-05 (CJO), Department of Health’s National Institute of Health and Care Research Biomedical Research Centre funding to University College London Hospital (JSB), Wellcome Investigator Award 221803/Z/20/Z (JSB), Medical Research Council Experimental Medicine grant APP11997 (JSB), NIH R01 grant AI114800 (CJO and HT), and the Universidad de La Sabana grant MED 285 2020 (LFR). We thank Karl A. Pruitt II, Katherine Kruckow, Daniel Minassian, and Chembilikandy Vipin for their assistance with *in vivo* work. We thank Ed Swords and David Briles for clinical bacteria isolates.

